# Cortical volume-to-surface and -to-white matter volume relations are explained by uniform cortical architecture in mammals

**DOI:** 10.1101/2020.08.16.252825

**Authors:** Marc H E de Lussanet, Kim J Boström, Heiko Wagner

## Abstract

The size of the mammalian cerebrum spans more than 5 orders of magnitude. The smallest cerebrums have a smooth (lissencephalic) cortical surface, which gets increasingly folded (gyrencephalic) with cerebral size. Further, the proportion of white-to-gray matter volume increases with the total volume. These scaling relations have unusually little variation. Even though a number of theories and models have been proposed, it remains an open question, why this is so. Here, we show that almost all variance is explained by assuming a homogeneous composition of the cortex across mammals. On the basis of this assumption we derive quantitative analytical computational models. The first model predicts the cortical surface area from the gray and white matter volume. A single free parameter, for the height of cortical columns is estimated as ***λ* = 2.9** mm (***r*^2^ = 0.996**). The second model predicts the white matter volume as a function of the gray volume and the cerebral size (with parameters for intra- and extra-gyral connections ***l_int_, l_ext_***; 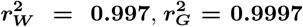). The models are validated by predicting the effective cortical thickness and the folding parameter ***κ***. The results accurately predict the human intraspecific variation of the surface relations. As expected, we find a reduced **λ** for cetaceans, and that preterm human infants do not follow the model. We also find deviations of gray and white matter volume for large cerebrums. Overall, the models thus show how the regular architecture of the cortex shapes the cerebrum. We conclude that the mammalian cerebrum scales in an *isomorphic*, rather than isometric, manner.

## 1 Introduction

Scaling relations are a classic field of evolutionary biology. The mammalian cerebrum ranges in size from about 10 mm^3^ (pygmy shrew) to more than 5 dm^3^ (fin whale, Pilleri and Gihr, 1970), i.e., more than 5 orders of magnitude. The scaling relations of the mammalian cerebrum are extraordinary because various parameters such as volume, surface, and proportion of gray matter (Changizi, 2001; Hofman, 1985; Cuntz et al, 2012) all show trends with very little deviation across several orders of magnitude and independently of phylogenetic relationships (e.g. Hofman, 1989; Zhang and Sejnowski, 2000). This is in contrast to other scaling relations such as the encephalisation (expressed as brain-, cerebral- or cortical mass to body volume) which can vary by twentyfold between species of the the same body mass (Boddy et al, 2012), and which have different scaling relations within species versus between species, and between phylogenetic groups (Boddy et al, 2012; Manger et al, 2012).

The tight scaling relations suggest a simple scaling mechanism that depends on only a few parameters. Understanding the common mechanisms that underlie the scaling relations will help to formulate scaling laws; to identify exceptions from these laws and thus help to understand evolutionary differences between species and brain regions (Rakic, 2009; Van Essen et al, 2019).

One remarkable scaling relation of the mammalian cerebrum exists between volume and total surface area (Hofman, 1985). Under the assumption that the overall shape of the cerebrum is scaleindependent (de Lussanet, 2015), this relation is equivalent to the relation between exposed surface area and total surface area, which is sometimes expressed as a ratio (called folding index or cephalization index: Manger, 2006). The cortex of small cerebrums (such as that of mice) has a smooth surface, which is called *lissencephalic.* The cortex of large cerebrums (such as that of cats, monkeys and humans) is increasingly folded, which is called *gyrencephalic.* Whereas isometric scaling of surface and volume exists for lissencephalic cerebrums, the total surface area of the cortex increases over-proportionately with the volume of the cerebrum (Braitenberg, 2001).

A second remarkable scaling relation has been found between the volume of gray and white matter (Zhang and Sejnowski, 2000). The authors also presented a theory for this relation which is based on an assumption of isometric scaling (i.e., that the grey matter volume scales isometrically with the average axonal length). Their model seems fits the grey-white matter volume relation moderately well, depending on the assumptions. In the appendix we prove that the isometric assumption is not tenable though.

According to one hypothesis the cortical folding is governed by a minimal free energy principle (Mota and Herculano-Houzel, 2015; Mota et al, 2019). This hypothesis was based on the so-called Flory-approximation for the folding statistics of long polymer chains in a solvent (Flory, 1979; Kantor, 2004; Nelson, 2004). Even though the hypothesis was published very prominently, it is wrong for two reasons. First, being a statistical approximation, the Flory model is only valid for very long chains or surfaces (where “long” is expressed in relation to thickness Kantor, 2004), but the surface of the mammalian cortex is not at all “long” in this sense (rather, the smallest, non-convoluted, cerebrums are as short as theoretically possible, whereas the largest cerebrums fit particularly bad to the model: de Lussanet, 2016). Second, the cerebrum is not a flexible surface whose folding is unrestrained by the surrounding “liquid” matrix. Rather, on the inside, the cortex is densely connected by white matter whose axons are to some degree under tension and thus exert a directed force on the cortical surface.

The above relations have been described for adult mammals, across species. Below we will show that the volume-surface relation is also valid within the human species. However, this does not mean that these relations are the same across embryological development. Indeed, this is unlikely because the cortical thickness increases dramatically during development while the relative volume of the ventricles decreases strongly.

According to an old idea, the convolutions develop because the cortex expands more strongly than the volume within (Le Gros Clark, 1945). Indeed, it has been shown that the swelling of gel that is stuck on a less elastic surface accurately mimics the structure of cortical gyri and sulci, and this was confirmed with a finite element model (Tallinen et al, 2016). Since swelling body expands not only in thickness (i.e., radially) but also in the tangential directions, its surface must expand, and if its base is restricted, it will spontaneously form gyri and sulci. According to another hypothesis, the tension of intracortical axonal connections might pull its surface into gyri (Van Essen, 1997; Essen, 2020). However, numerical simulations and dynamic tensor imaging (DTI) failed to confirm this idea (Xu et al, 2010).

An alternative tension hypothesis has proposed that the axons that are oriented perpendicular to the cortical surface evoke a buckling pattern of the cortical surface, especially in large cerebrums. This hypothesis is supported by a numerical computational model (Manyuhina et al, 2014). Indeed, local changes in the main orientation of axon fibres (as shown by local minima in fractional anisotropy in DTI (diffusion tensor imaging) of sheep embryos seem to coincide with the future location of the primary sulci (Geng et al, 2009). Accordingly, the primary sulci, which develop first, are the deepest and most consistent across species, are the ones that coincide with the primary cortical sensory and motor areas (de Lussanet, 2021).

It is widely accepted that the mammalian cerebral cortex shows a highly homogeneous composition of layers and neuronal cortical columns. The layers differ between regions of the mature cortex, but always show the developmental state of six layers at some time (Brodmann, 1913). Cortical columns integrate the layers perpendicularly to the cortical surface, and are regarded as computational units, which are highly uniform across the size range of the cerebrum (Rakic, 1995). It is thought that the volume of cortical columns is independent of the local curvature of the cortical surface (Bok, 1929). Evidence from precise counting studies across species suggest that the number of cortical neurons per square millimeter is invariably close to ~10^5^ per mm^2^ (Rockel et al, 1980; Carlo and Stevens, 2013).

White matter contains the myelinated longdistance axonal connections. To answer the question how the volume of the white matter relates to the gray matter it is, therefore, essential to know how the diameter of axonal connections depends on the length of the axons. For example, it is well known that the myelin sheath increases the signal transmission velocity linearly with the axonal radius (Perge et al, 2009). Many invertebrates possess a few giant axons for long-range startle responses. Accordingly, one might expect large mammalian cerebrums to have axons with a wide diameter.

However, since the axonal volume increases quadratically with its diameter, a bundle of many narrow axons will transmit more information per second than a bundle with the same cross-section, which consists of a few wide axons. Moreover, due to noise, the efficiency of information transfer decreases with the spike rate according to Shannon’s information theory (Balasubramanian et al, 2001). Indeed, it has been estimated from the frog’s retina that each cell type transmits approximately the same amount of information independent of its axonal diameter, so that thin fibres transmit vastly more information per summed cross-sectional area than thick ones (Koch et al, 2006). Consistently, the axonal width distribution in the mammalian brain seems to be largely independent of brain size (Perge et al, 2012).

In the present study we aim to explain the empirical scaling relations of the mammalian cerebrum. We hypothesise that valid scaling laws can be formulated on the basis of a simple growth rule and the uniform structure of the cortical building blocks and the scale-independent distribution of axonal widths. Below we derive a model that predicts the total cortical surface area on the basis of the grey and white matter volume. One parameter is fitted, i.e., the average height of the cortical columns. We hypothesize that a single scaling relation holds for the entire size range of mammalian cerebrums, independent of phylogenetic relations. The model is fitted to existing empirical data of the cerebral volume to surface area.

Similarly, to calculate the white matter volume, we hypothesize that the distribution of axon diameters in the white matter is independent of cerebral size and that the number of axons scales linearly with the grey matter volume. This second model with two free parameters will be fitted to existing data of cerebral grey and white matter volumes.

The models will be validated using existing empirical data of 1st the exposed surface area to total surface area, 2nd we hypothesize that the same scaling relation holds within a species, i.e., the human cerebrum, 3rd the ratio of grey matter volume and surface area to cerebral volume (*effective thickness*) and 4th the folding parameter (*κ*).

Third, having developed a theory-based and validated tool allows to investigate exceptions. The tension hypotheses are developmental ones, and so are predicted to hold even for the growing cerebrum after birth. However, before birth, there is no need for functional cerebral sensorimotor control and so it is expected that preterm infants deviate from the predicted relationships. Also, we will discuss the results for cetaceans in the light of earlier results.

## 2 Modelling

### 2.1 Surface to volume scaling relation

This model will result in two equations, one for convoluted or *gyrencephalic* cerebrums (Eq. [7]) and one for smooth or *lissencephalic* cerebrums (Eq. [8]). The predicted relation is the maximum of these relations.

The outer surface (a.k.a. “exposed” surface) of the cerebrum differs from the deeper parts: 1. it does not contain any white matter, only gray cortex; 2. it is formed of the apex of gyri, so that the cortex in this border region is more strongly curved than the cortex inside the sulci.

We can thus distinguish the cortex (with volume *G*) into a convex gyral border of volume *G_B_* and thickness *λ*, and a planar to concave sulcal core of volume *G_C_* and a radius *R* – *λ*. Since the fundus of the sulcus occupies a very small part of the total cortical surface, it can be neglected and it follows that the area of the core surface equals

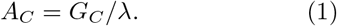

The volume *V_C_* of the core consists of gray and white matter, so we have *V_C_* = *G_C_* + *W*, with *W* from Equation [11]. Simultaneously, the core volume is approximately equal to the volume of a ball of radius *R* – *λ* (see above), so

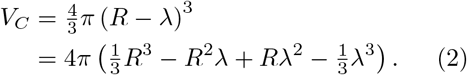

Using Eq. [1] and [2], the core surface area *A_C_* can thus be expressed as

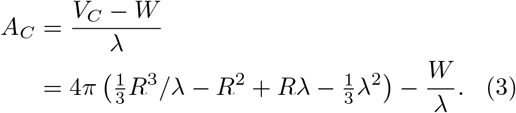

The cerebral border region *V_B_* is free of white matter but composed entirely of the convex part of the gyri. Due to the convexity, the cortical columns are not column-shaped as in the sulcal regions of the cortex, but rather pie-shaped. Assuming that the cortical columns in the convex regions have the same volume as those in the flat, sulcal regions, the cortical surface per cortical column should be about twice as large in the cerebral border region. For the cortical surface in the border region *A_B_* that:

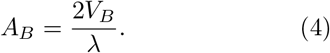

The border volume is the cerebral volume minus the core volume, hence *V_B_* = *V* – *V_C_*. Let us approximate the cortical volume *V* by the volume of a ball of radius *R*, hence neglecting the sphericity, so 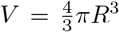, and let us use Eq. [2] for the core volume *V_C_*, we then arrive at

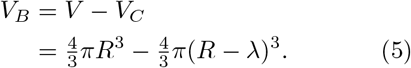

Inserting the above expression for *V_B_* into Eq. [4], we obtain

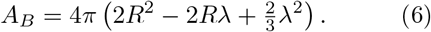

Using the above equation and Eq. [3], the total cortical surface of a gyrencephalic cerebrum reads

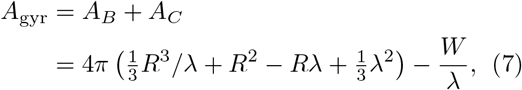

where 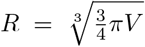 (because 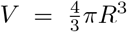 as explained above).

If the radius of the cerebrum is smaller than the thickness of the border, the cerebrum is smooth (lissencephalic). In that case, the cortical surface equals the spherical volume and surface relations (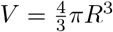 and *A* = 4*π R*^2^), so

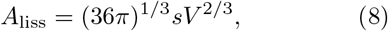

with the dimensionless sphericity constant *s* = 1.56 to correct for the deviation from a perfect sphere of the cortical hemispheres (Fig. S1).

Equations [7] and [8] thus predict the scaling relation for the cortical surface. The intersection point between the two equations is the transition between them, such that

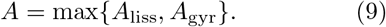

The white matter volume *W* (eqn. [7]) can, unfortunately, not be obtained directly from published data. Instead, the white matter volume will be estimated using eqn. [11] below. The model has thus just one single free parameter λ to be fitted against the data.

### 2.2 Gray and white matter volumes

Assuming that the distribution of axonal diameters in the white matter is independent of brain size (Perge et al, 2012, see introduction), the volume of the axonal connections increases linearly with the distance between the regions that they connect. The summed cross-section of the axonal connections scales with the number of neurons in the gray matter, which in turn scales with the gray matter volume *G*. Since the white matter volume *W* is proportional to the mean axonal length 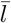 times the summed cross section, it follows that the white matter volume is proportional to the average length of the axonal connections (i.e., 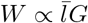, where length parameter 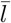 is dimensionless).

The 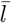 is a weighed sum of local, intra-gyral, connections and extra-gyral ones, given our central assumption of a scale-independent composition of the cortex. The length of intra-gyral connections is independent on the cerebral size, and is thus a dimensionless constant, *l_int_*.

The length of long-distance extra-gyral connections scales with the radius *R* of the cerebrum. Given the characteristic overall shape of the cerebrum, the total volume of the cerebrum scales proportionally with the cube of the radius, so that we obtain

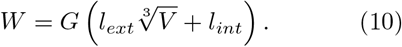

with *l_ext_* the dimensionless length constant for extra-gyral connections.

Since *V* = *W* + *G* we obtain for the gray and white matter volume:

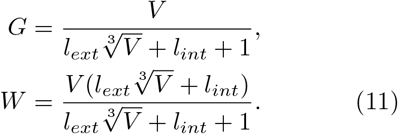

## 3 Results

The model for the relation of cortical surface to cerebral volume has just one single free parameter *λ*, which describes the cortical thickness. Fitting this parameter to the averages per species (excluding the human MRI data) and using a mean sphericity of *s* = 1.57 (Fig. S1), resulted in *λ* = 2.81 mm (N=47, *r*^2^ = 0.996; Fig. 1A). (The cetacean λ= 1.9. Without the cetaceans, the fit would be even slightly better: *λ* = 2.94 mm, *r*^2^ = 0.997.) The residuals (log units) are shown in Panel B (curvature: P < .001). The fitting of the individual measurements in the human dataset resulted in *λ* = 2.97 mm (*r*^2^ = 0.962; Panel A-inset). The corresponding residual is shown in the inset, curvature: *P* < .01. For a part of the measurements (N=51 species) the exposed cerebral surface area was reported. Using *σ*, the cerebral volume was estimated for these measurements (Fig. 1C). The residuals (Panel D) systematically increase with cerebral volume (*P* < .01). Note, that the lissencephalic cerebrums are exactly on the predicted relation because cortical and exposed surface area are equal.

**Fig. 1.**
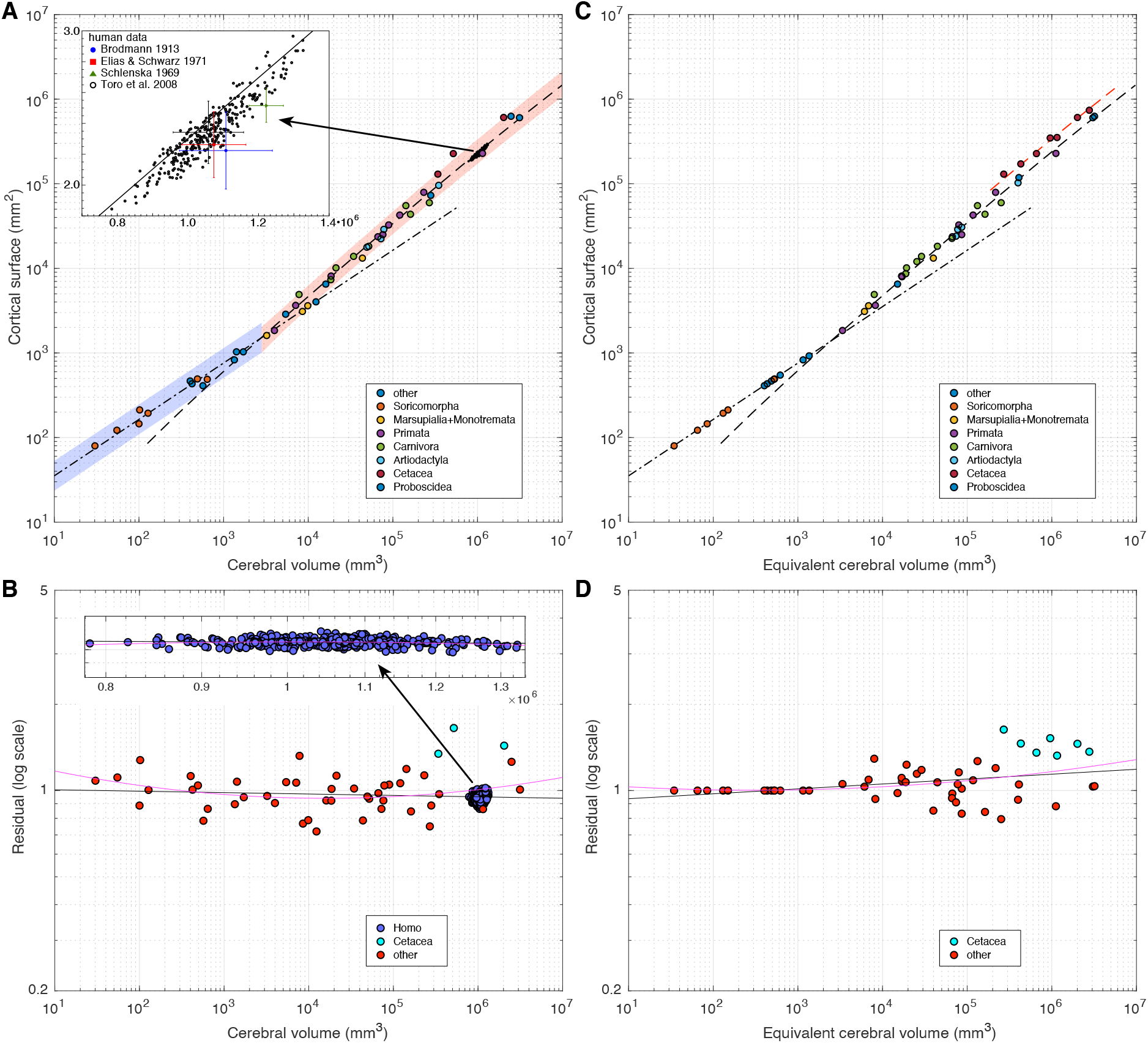
Results of model 3 for the cortical surface in relation to the cerebral volume. A. Gyrencephalic cerebrums scale with Eq. [7] (dashed curve); lissencephalic cerebrums scale with Eq. [8] (dash-dotted curve). Shaded regions show a factor 1.5 for the sphericity, *s* (blue) and cortical thickness *λ* (red) parameters. Note that the set of human MRI data (cloud of dots), was not included for the data fitting. Inset: The human data by literature source. B. The residual for panel A on log scale as a function of the cerebral volume. Black lane and lilac curve: linear and quadratic regression, respectively (including the cetaceans, blue dots). The inset shows the human MRI data with the same vertical scale. C. The same model; the equivalent volume was calculated from the exposed surface area, using the exposed surface to volume ratio (*A_exp_*, see Fig. S1 and Methods section). The black curves are the same as in Panel A. The red dashed curve shows the model fitted to just the cetaceans. D. The residual for panel B. Note, that the exposed surface equals cortical surface for lissencephalic cerebrums. The cetaceans are shown with blue dots.

Model 2. The gray and white matter volume model, Eq. [11], describes the data very well, with the best-fitting parameter values *l_ext_* = 0.00811, *l_int_* = 2.060 (with 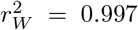; 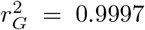). The resulting two relations are plotted in Figure 2A. The residuals (log units) were generally much smaller for gray then for white matter volume (Fig. 2B: gray slope: *P* < .05, curvature: *P* < .01; white curvature: *P* < .05).

**Fig. 2.**
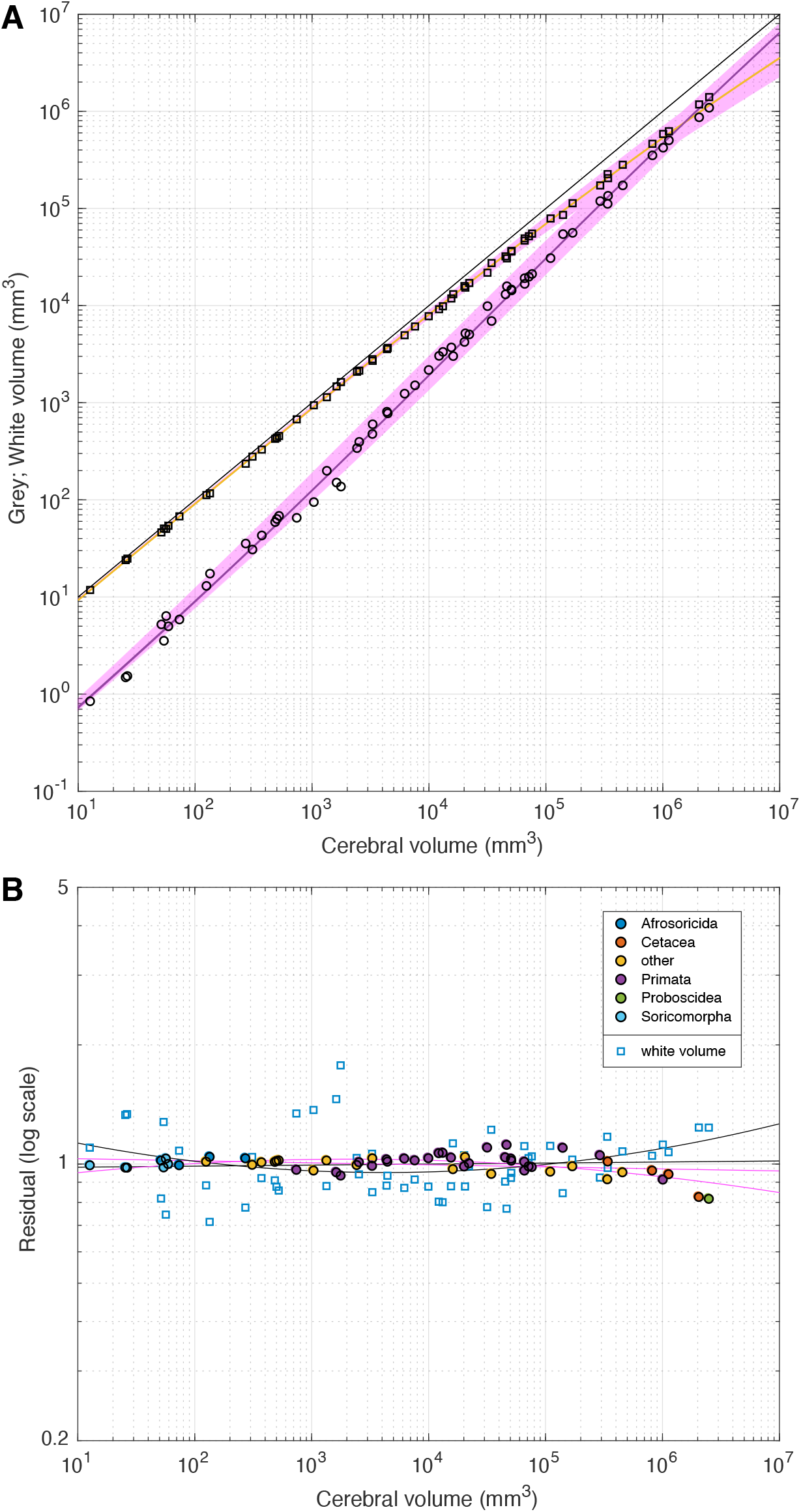
Model for gray and white matter volumes, fitted to the dataset of Zhang and Sejnowski (2000). According to Model 2 (Eq. [11]), the white matter volume scales to the cerebral size in two ways: long-range extra-gyral connections scale with the cerebral radius, and short-range, intra-gyral connections scale with the total cerebral surface. A. The fitted relation on a log-log scale. B. The residual on a log scale as a function of the cerebral volume. Note, that the residual has log-units. Black curves: linear and quadratic regressions to the white matter residual; purple curves: the same for the gray matter.

Two existing relations can be predicted on the basis of the current models. The *effective cortical thickness* is defined as the ratio of estimated gray matter volume (eqn. [11]) to cortical surface area (eqn. [7]) represents (cf. de Lussanet, 2016), and is shown in Fig. 3A. The model (red dashed curve) describes the general trend in the empirical data, but the data have considerable variation for gyrencephalic cerebrums.

**Fig. 3.**
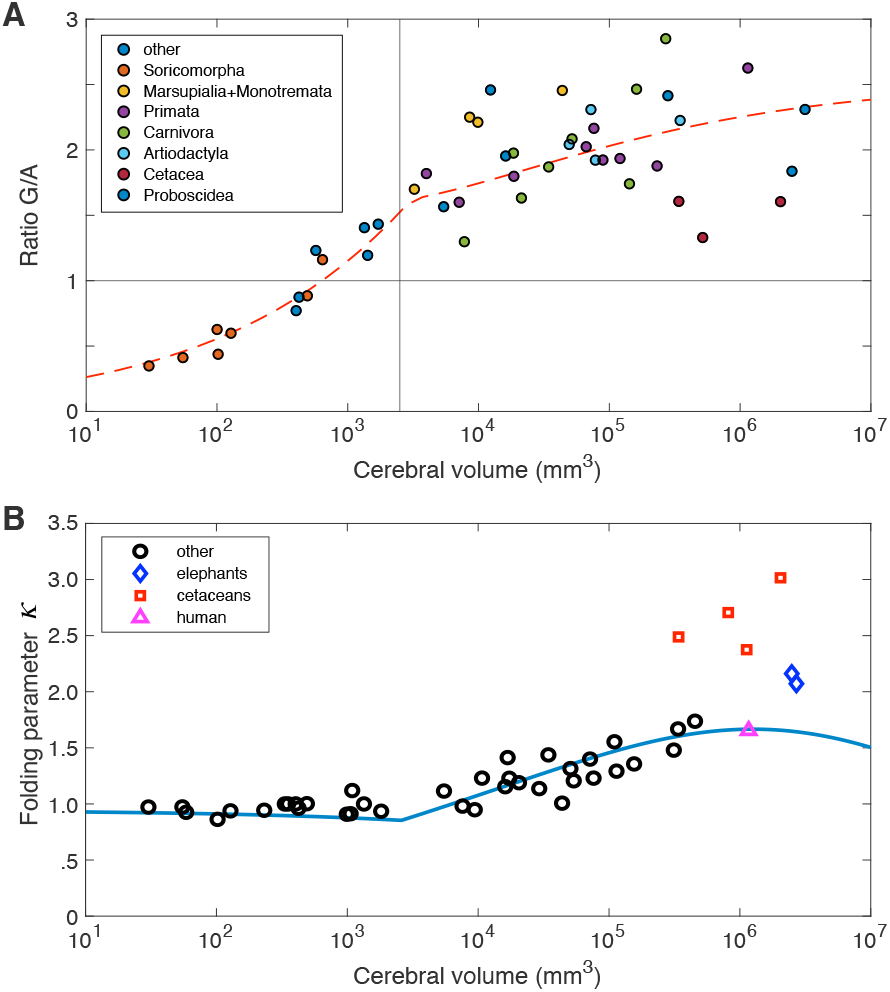
Predictions for the effective cortical thickness (A) and folding parameter *κ* (B). A. The effective cortical thickness, i.e., the ratio of gray volume to cortical surface area *G/A* as a function of the cerebral volume (log(*V*)). Vertical line: transition to convoluted cerebrums. Dashed curve: Model prediction (see text). B, The folding parameter *κ* is the proportion of grey volume times the folding index (de Lussanet, 2016), as calculated from the models along with the data from (Mota and Herculano-Houzel, 2015). Blue squares: cetaceans, red rhombus: African elephants.

The folding parameter *κ* was predicted to be a constant by the model of (Mota and Herculano-Houzel, 2015), but this prediction was falsified on a statistical basis (de Lussanet, 2016). The folding parameter can also be predicted on the basis of the equations [11] and [7] and the spphericity (see Supplement). The prediction, along with the empirical data (cf. Mota and Herculano-Houzel, 2015), are shown in Figure 3B.

## 4 Discussion

We have proposed a unified theoretical concept to explain the scaling relations of white matter volume and cerebral surface area. The basis of the concept is simple and can be addressed as *isomorphic scaling law*: the regular scaling patterns of the mammalian cerebrum are caused by, and reflect, the highly regular and uniform composition of the cortex into a fixed number of (six) layers and into a variable number of cortical columns. The presented models are derived on the basis of these premises.

This isomorphic scaling concept differs clearly from the well-known classical isometric scaling concept and does not lead to the well-known isometric scaling relations. However, the simple and clear underlying principle nevertheless leads to remarkably narrow empirical scaling relations that are very well described by simple analytic scaling laws (equations 9 and 10).

The first model (Eq. [9]), contains just one single free parameter: the cortical thickness *λ*, while fitting the data really well (*r*^2^ = 0.996). Across species the cortical thickness (*λ* = 2.81 mm) was very close to the intraspecific value for humans (*λ* = 2.97 mm). This latter value is within the range reported by Bok for a single specimen (Bok, 1929). Our value for *λ* is somewhat higher than the *effective* cortical thickness (i.e., the ratio of *G/A* = 2.63 for humans, cf. Fig. 3A; see also Fran-gou et al, 2020). This is to be expected because the latter definition neglects that the cortical surface has a convoluted shape which is overall convex.

As argued in the introduction, the sulci are of varying depth and so show a hierarchy (Garcia et al, 2018). This hierarchical folding pattern can be formulated as rule leading to a very simple model describes the scaling of the volume-surface relation quite well (Fig. S2). Since this model also follows the isomorphic assumption, it is consistent with our hypothesis despite some crude simplifications (it neglects the white-matter volume and is only quasi 3-D). An interesting aspect, however, is that it implements a fractal-like shape rule. This notion has existed since several decades (Hofman, 1991; Mota and Herculano-Houzel, 2015; Stoop et al, 2013), but could not be demonstrated yet (de Lussanet, 2016).

The second model derives the white matter volume by the additional assumption that the cross-section of white matter per gray matter volume is constant, and that a constant proportion of the white matter connections are connecting locally versus globally. With just two free parameters, the model fits the data well (*r*^2^ equals 0.997 and 0.9997, respectively, see results). The small quadratic trend in the residual suggests that the model slightly overestimates the white matter volume in the smallest and the largest cerebrums. For small cerebrums this is indeed to be expected, when the radius of the cerebrum becomes smaller than the length of the local connections in a gyrencephalic cerebrum. For elephants and pilot whales (the two data points with the largest cerebrum *V* ≈2 l) the model predicts the white matter volume to be larger than the gray matter volume, which is not the case. Rather, the proportion of white matter seems to saturate around 45% for brains the size of humans and larger.

Zhang and Sejnowski (2000) proposed a model for the grey-to-white matter volume relation on the assumption that the cortical geometry minimizes the overall length of the axonal connections (maximizing compactness), predicting that *W* ∝ *G*^4/3^ (see Introduction). The power of scaling is represented by the slope in a log-log graph. Linear regression on the log-log data resulted in a significantly lower slope of 1.23. By contrast, our model predicts the scaling power well, increasing slightly with cerebral size (from 1.09-1.36).

The folding parameter was proposed by (Mota and Herculano-Houzel, 2015) as a fitting constant of their model assuming optimal free energy in tension for the cortical surface area (see Introduction). de Lussanet (2016) showed that this parameter is better expressed in terms of the proportion of gray matter volume and the folding index: *κ* = *G/V Ȧ*/*A_exp_*. The latter study showed that *κ* deviates significantly with cerebral size. Here we show, that the folding parameter *κ* is well predicted by the current models (Fig. 3B), with the exception of cetaceans and elephants, due to the saturation in the grey-white matter volume relation (see above). Thus, the folding parameter provides additional evidence, validating the current modelling approach.

A chronic difficulty of earlier models has been to predict the power of the scaling relations for the cortical surface (Hofman, 1985, 1989; Mota and Herculano-Houzel, 2015; Xu et al, 2010; Braitenberg, 2001) as well as for the white matter volume (Zhang and Sejnowski, 2000). Hofman 1989 found a slope of 0.83 for a linear regression on convoluted cerebrums. The current model predicts a power of scaling is very close to 0.83 within the size range of 10^4^ – 10^7^mm^3^ and a slightly higher power for smaller sizes.

The reasoning underlying the present scaling theory is that across the huge size range, deviations from the general trend are small. It follows that few factors determine the general scaling pattern, so we derived analytic scaling relations. Obviously, this approach involves *simplifications.* For example, the number of neurons (Rockel et al,1980; Carlo and Stevens, 2013) and the effective cortical thickness (Frangou et al, 2020) are slightly different between the different lobes. However, since such local differences are present across species of different sizes (Rockel et al, 1980), they will average out. Also, the cortical thickness *λ* depends on the curvedness (Bok, 1929). The model will therefore slightly overestimate the cortical surface area in the border region. On the other hand, it will slightly underestimate the cortical surface in the core region because the core region also contains gyri. The central conclusion is, that, indeed, simple principles can be defined to predict almost the entire variation in white matter volume, cerebral surface area and folding pattern.

One might speculate about deviations (and the absence thereof) with respect to the models. We found no systematic effects of different literature sources despite the data being collected across an entire century using various methods (cf. supplementary Fig. S3) B). One important source of error might be in the conservation artifacts (all studies corrected for those artifacts). The small systematic differences in cortical surface area in human data from conserved specimen (Brodmann, 1913; Elias and Schwartz, 1971) with respect to the in-vivo MRI data (Toro et al, 2008) might be due to the older age of the subjects in the earlier studies, which were analysed post mortem (Fig. 1A, inset). The overall slightly larger volume of Schlenska (1969, N=5) might be due to the small sample and the conservation artifacts. Given that the standard deviations in all four studies are very similar, suggests that the precision of the techniques used is good and relatively small in comparison to the natural variation in cerebral size.

Another remarkable pattern is the lack of phylogenetic trends. In cerebrums larger than human’ s, the relative contribution of gray and white matter seems to saturate (Fig. 2B). This observation involves the African elephants and cetaceans, and is clearly reflected in the folding parameter *κ* which the current model even predicts even to go down for very large cerebral sizes. This deviation probably reflects that the assumption that the long-distance connections scale with cerebral radius (eq. [10]), does not hold for very large cerebrums. More data are needed to test this.

The only clade that shows systematic deviations from the cortical surface model (Fig. 1) are cetaceans (Manger et al, 2012; Roth, 2015; Marino et al, 2007). The data strongly suggest that cetaceans have a lower cortical thickness parameter *λ* (cf. Fig. 1C). This is consistent with 40% increased cortical surface area with respect to the entire cerebral volume. For example the number of glia cells, e.g. astrocytes, throughout the cerebrum could be enhanced. Indeed, evidence shows that the large cerebrum cetaceans (and other aquatic mammals) does not provide them with superb cognition or intelligence, a lack of neuronal specialization and prefrontal cortex (Manger, 2006, 2013; Patzke et al, 2015). Rather, the increased brain size seems to reflect a physiological specialisation. Manger (2006) argued that this physiological specialisation is heat production: the mammalian is highly sensitive to minor changes in temperature, so keeping brain warm in cold water should indeed present a major evolutionary pressure.

Humans, on the other hand, are very close to the model fit for both models. This finding is in line with earlier conclusions (Azevedo et al, 2009; Herculano-Houzel, 2012). The data of Toro et al. (Toro et al, 2008) show, moreover, that the model predicts the scaling relation even within the human species well for young adult humans (Fig. 1B).

We hypothesized that preterm infants will be below the predicted scaling relation because the cortical thickness decreases with growth. The MRI data of prematurely born infants (Kapellou et al, 2006) suggests that the predicted scaling relation is reached around the age of normal gestation time (38-42 weeks: Fig. 4). According to these data, the cerebrum is lissencephalic until about 10^5^ mm, i.e., 50 times the transition to gyrencephaly in adult mammals (Fig. 4). These data thus suggest, that the scaling relation might be accurately predicted across the developmental span for humans post gestation. This is consistent with Wang et al (2017) who found that the increase in cortical surface area comes later than the increase in cortical volume.

**Fig. 4.**
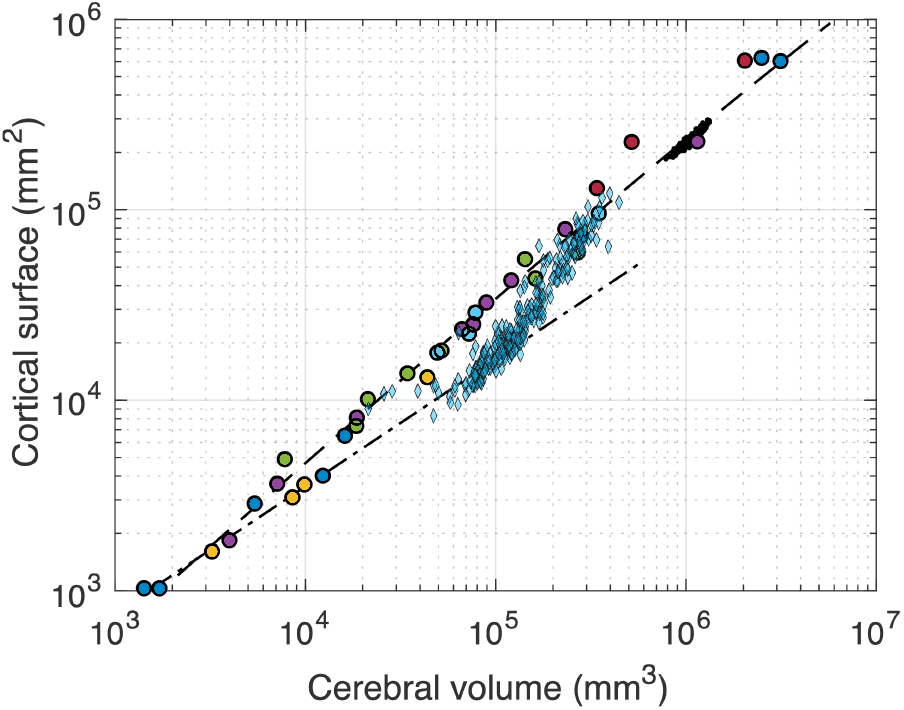
Very premature infants scale differently. The figure shows an inset of Figure 1, with additionally included MRI data of 130 extremely prematurely born infants (blue triangles; gestation week 22-29; normal full term would be 38-42 weeks;Kapellou et al, 2006). Most infants were scanned more than once (n=274). Age range at scan: 23-48 weeks post gestation. The volume-surface relation was much more variable than for the adult population. The smallest cerebrums were lissencephalic until about 10^5^mm, i.e., 50 times the transition to gyrencephaly in adult mammals. From there, a steep increase in cortical surface occurs (the black line represents *A* = *V*^2^). The model curve is reached at a volume of about 2 – 4 · 10^5^mm, which is only about 1/3 of the average adult volume.

A considerable number of models have been proposed to explain the cortical folding and the scaling relations of the mammalian cerebrum. Most have not been validated to empirical data (e.g. Karbowski, 2003; Schlenska, 1974; Jerison, 1979; Hofman, 1985, 1989; Prothero and Sundsten, 1984; Todd, 1982; da Costa Campos et al, 2020; Van Essen, 1997; Mota and Herculano-Houzel, 2012). Other models were validated (e.g. Mota and Herculano-Houzel, 2015; Xu et al, 2010; Braitenberg, 2001; Zhang and Sejnowski, 2000), but failed to explain the empirical data. This state of research is confusing, given that the empirical data show such an unusually clear pattern, which suggests that the scaling relations are governed by just few parameters.

We have here proposed two models that explain the pattern of cerebral folding (gyrification) and white matter volume, on the basis of a single concept, i.e., the homogeneous construction principles. According to this principle, both white matter volume and cortical surface area are simple geometric scaling functions of the gray matter volume. The validation to the available empirical data shows that model and data are fully consistent.

### Conclusions

1. On the basis of very few simple principles it is possible to predict almost the entire variation in white matter volume, cerebral surface area and folding pattern.
2. Our model is validated by the available empirical data.
3. Our new theory provides strong evidence that uniform cortical composition in mammals drives the most important scaling relations known so-far as an isomorphic scaling law.
4. We provide evidence that the cortical surface area and the volume of white matter in the mammalian cortex are governed by the volume of gray matter, probably due the cortex being universally composed of cortical columns and being layered.
5. The hierarchical and evolutionary conserved pattern of sulci, formulated as an isomorphic fractal-like growth rule describes the scaling law well.
6. The proposed scaling relation is largely independent of phylogenetic relations and even holds within the human species.
7. According to the present view, the regular scaling relations of the cerebrum do not reflect an evolutionary optimum. Rather, the present scaling laws can be formulated on the basis a simple growth rule and the uniform structure of the cortical building blocks.

## 5 Methods

### 5.1 Datasets

For the gray and white matter volume data, we used the database of Zhang and Sejnowski (2000), which includes measurements for 59 species of mammals. The cerebral volume and cortical surface area for human cerebrums was measured for 314 young adults on the basis of in-vivo MRI measurements (Toro et al, 2008).

For the database of surface to volume, only measurements from the same specimen were included (see Supplementary Table). If for a species several measurements were available, these were averaged. Since not all studies measured all parameters, the number of included measurements may differ depending on the included measures. For example: three studies report data for horses, in total 7 specimen. When including cerebral volume and total surface area the mean volume for the horse is 297 cm^3^ (N=7), but when including cerebral volume and exposed surface area, the mean volume is 450 cm^3^ (N=2), because the exposed surface area was not measured (Mayhew et al, 1996).

The first systematic and accurate measurements of a wide range of mammals have been developed and published by Brodmann and Henneberg (Brodmann, 1913; Henneberg-Neubabelsberg, 1910). The total cortical surface was measured by precisely covering it with pieces of silk paper. Since this method is not easy to apply and is very labour intensive, later measurements tended to use a statistical approach using slides, e.g. counting the number of surface crossings on a grid and applying a correction formula. A further problem is the effect of shrinkage due to conservation in ethanol (e.g. Mayhew et al, 1996). Consequently, it is likely that an important source of variability in the data lies is in the measurement techniques used.

The selection criteria for the inclusion of studies were as follows: 1. Systematic usage of a validated method; 2. Reporting of a range of mammals (not just humans); 3. Reporting of at least the cerebral surface area and the cerebral volume and/or the cerebral outer surface area from the same specimen. Seven studies matched these criteria: (Brodmann, 1913; Schlenska, 1969; Haug, 1970; Elias and Schwartz, 1971; Schlenska, 1974; Mayhew et al, 1996). Finally, the measurements of reported total/outer surface ratio of 1.0 (i.e., lissencephalic cerebrums) were added from Hofman (1985).

Exclusion criteria were applied as follows (see Supplementary Table): 1. Malformed or diseased brains (this applies to some measurements in Brodmann, 1913); 2. excessively deformed brains (Brodmann suspected excessive shrinkage of two brains of Ganese origin); 3. Estimates (e.g., of cerebral volume) that were based on other specimens or taken from the literature; 4. Measures of cerebral volume that do not fit into a sphere of the reported total cortical surface (this applied toElias and Schwartz, 1971); 5. for *Homo sapiens,* ten statistical outliers from three publications were excluded.

The remaining dataset used in the present study thus includes 141 samples (incl. 29 humans) from 55 species and 17 orders of mammals. The cerebral volume was available in 97 samples and the outer surface of 104 samples. For the fitting a single average per species was calculated: 44 species for the *A_c_*-*V_c_* fit and 51 species for the *A_c_*-*A_exp_* fit. In either case, the *A_c_* of a species was calculated over the samples that were available for the second parameter, so that the mean *A_c_* for a given species could differ depending on the fit. The complete data set as well as a commented listing of the excluded measurements is provided for downloading (see Supplementary Table).

### 5.2 Analysis

The fitting procedures were performed on the mean values of the species, so that each species is represented by just a single value regardless of the number of samples that had been used for the mean. These data were fitted using a least squares fit to the logarithmic data. To estimate the goodness of fit, not only the r^2^ were taken, but also the first and second order trends in the residue. Such linear or u-shaped systematic deviations from the model can be important cues for validating the underlying model.

The *Supplementary Excel Table* presents the data collected from existing literature sources as well as the reasons why specific measurements were excluded. Further, the calculation of the means is shown in an extra sheet.

## 6 Declarations

The authors have no competing interests to declare that are relevant to the content of this article. No funding was received for conducting this study.

### 6.1 Author Contributions

MdL designed and performed the research, analyzed the data, wrote and revised the critically manuscript. KB and HW designed the research and critically revised the manuscript. All authors gave final approval for publication and agree to be held accountable for the work performed therein.

## Supplement

### Dataset

The supplementary excel table presents the data collected from existing literature sources as well as a list of notes for the reasons why specific measurements were excluded. Further, the calculation of the means for each species is shown in an extra sheet.

### Overall shape of cerebrum

For several considerations the relation between the volume *V* and the “exposed” surface *A_exp_* (i.e. without the sulcal surface) of the cerebrum is of relevance. If the overall shape is constant, the ratio *A_exp_*/*V*^2/3^ should be constant. This is shown in Figure S1. See also (de Lussanet, 2015).

**Fig. S1.**
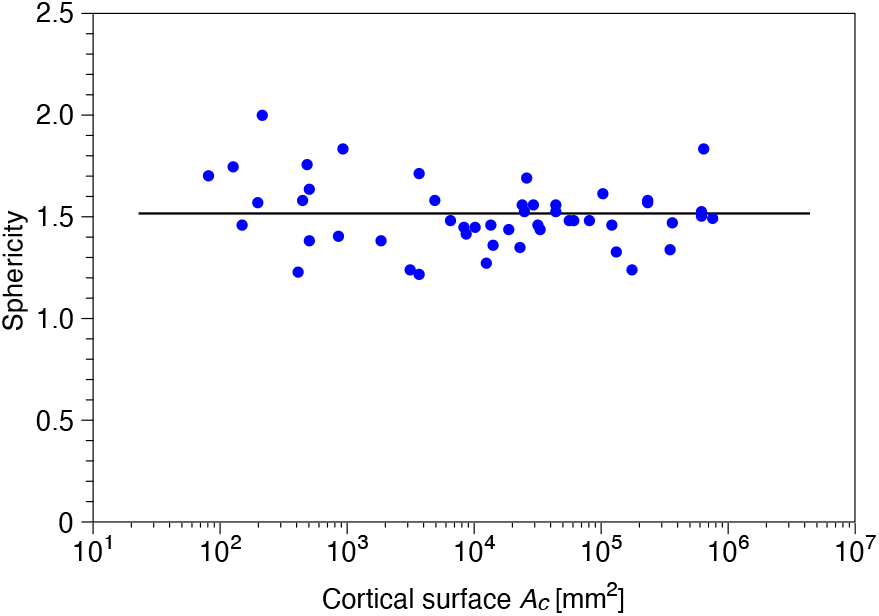
Sphericity of a 3D shape is defined as the ratio of the exposed surface (*A_exp_*) to the surface area of a sphere expressed in terms of its volume: 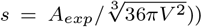. The equation for the surface area of a sphere of volume *V* can be calculated by solving the standard equations for it surface area (*A_sphere_* = 4*π R*^2^) and volume (*V*_sphere_ = 4/3*π R*^3^). According to the database, the mean *s* = 1.57 ±0.16 (st.dev.) for the mammalian cerebrum (N=43 species, 66 specimen; see supplementary Excel Table).

### Grey-white matter relation is not isometric

To our knowledge, two hypotheses have aimed to explain the scaling relation between grey (G) and white (W) matter volume for the mammalian cerebrum (Zhang and Sejnowski, 2000). These authors proposed that

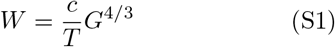

where *c* is a constant and *T* the cortical thickness (in the current work referred to as *effective thickness*). They argue that *T* is proportionate with *G*^0.1^ even though this is not well supported by the data (e.g. Hofman, 1988). Since *T* refers to the effective thickness, *T* = *G*/*A*, their equation be written as

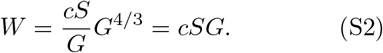

This form is not only simpler, but also more consistent with the data that have typically been reported in the literature (see supplementary Table).

### First order approximation: Fractal-like growth rule for gyrification

The first order model neglects the white matter contribution to the cerebral volume. It is based on the central hypothesis of a homogeneous composition of the cortex. Accordingly, the thickness of the cortex is limited, so once the critical thickness is reached, growth is only possible if the cortex invaginates, to form a sulcus (Fig. S2A). If the cerebrum exceeds a critical size (inner section “1” in Fig. S2A), a sulcus is initiated, but as long as it is smaller than the critical size, its surface is smooth (lissencephalic).

According to the first model, growth only occurs on the apex of each gyrus, and so the valley of a sulcus, once formed, will not move away from the center. Thus, the number of sulci decreases linearly when moving from the surface to the center of the cerebrum.

Note, that for very small cerebrums, the surface is entirely on the outside, so the cortical surface *A* scales to the cerebral volume *V* according to an isometric scaling law: *A* = *cV* (with *c* being a constant). On the other hand, for very large cerebrums, most of the cortical surface lies within the sulci, and since the thickness of the gyri remains constant, the surface will scale to the volume in a linear manner. Thus, the model will interpolate between the two scaling laws: a 2/3 power law for very small cerebrums and a linear scaling for very large ones (cf. Fig. S2B).

Note that this simple, first order model is just two-dimensional (2D), which is sufficiently realistic, though, because primary fields are elongated, so that gyri and sulci are elongated as well.

Analytically, the total circumference *C* of the 2D model is the sum of the exposed and nonexposed (inward) circumferences, which can be calculated as: 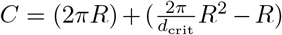 for a circle with radius *R* and straight sulci. Additionally, a shape constant can be introduced to account for the deviation of an exact circular shape of the model, the 3D version of the model will thus be of the general form: *C* = *R*^2^/*d* + *sR*, where *d* has the dimension of length and is related to the critical distance *d_crit_,* and where *s* is a dimensionless shape parameter.

When adding the third dimension, the total circumference becomes the total cortical surface A. The exposed surface will scale with the volume *V* to the 2/3 power in analogy to the surface of a sphere, so *V* ~ *R*^3^, whereas the non-exposed, inward folded surface scales again linearly with the volume (as a result of the invariant gyral thickness), so *V* ~ *A*^3/2^. In summary, the 3D version of the model is of the general form *V* = *A*^3/2^/*d* + *sA*, where *d* is, again, a thickness parameter with the dimension of length, and where *s* is *a* dimensionless shape parameter. Note that d and s in the 2D and 3D models are different but analogous parameters.

**Fig. S2.**
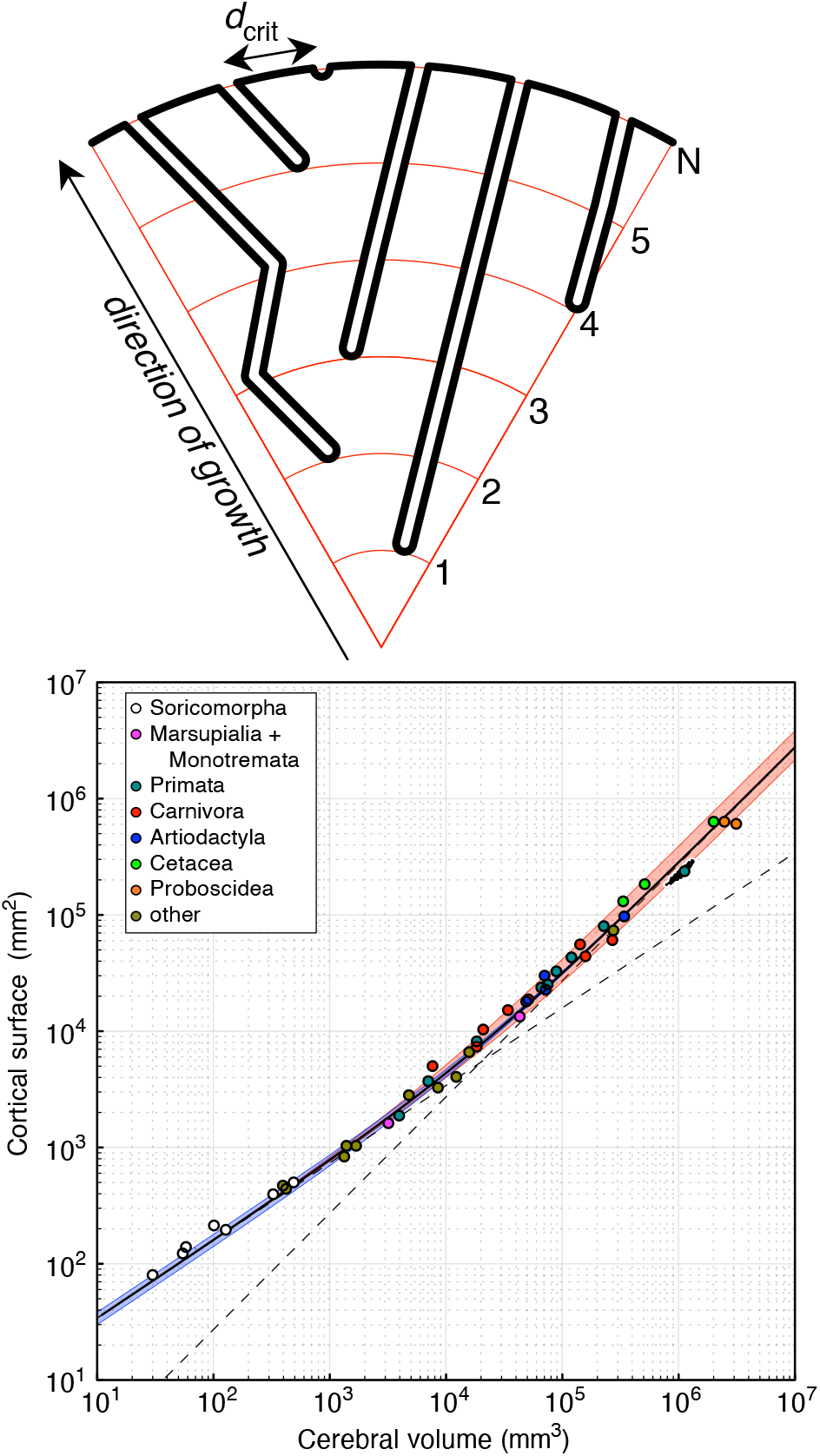
A. Schema of fractal-like growth rule approximation: The cortical surface (thick black curve) is on the exposed outside as well as in the sulci. Each time the gyral width exceeds the critical width (*d_crit_*), a new sulcus is formed. By this rule, the number of sulci, *N*, increases with the radius of the cerebrum. Note, that a small cerebrum with not have any sulci and will thus be lissencephalic (smooth). B. Behavior of this first order approximation on a logarithmic scale. Each dot represents the mean of one species (human data are in addition shown as a cloud). On a logarithmic scale, the model approaches two asymptotes: *A* = 7.1V^2/3^ (isometric scaling relation), *A* = 0.22*V* (linear relation). Blue and red shades: influence of 1 standard deviation for each of the two fitting parameters.

**Fig. S3.**
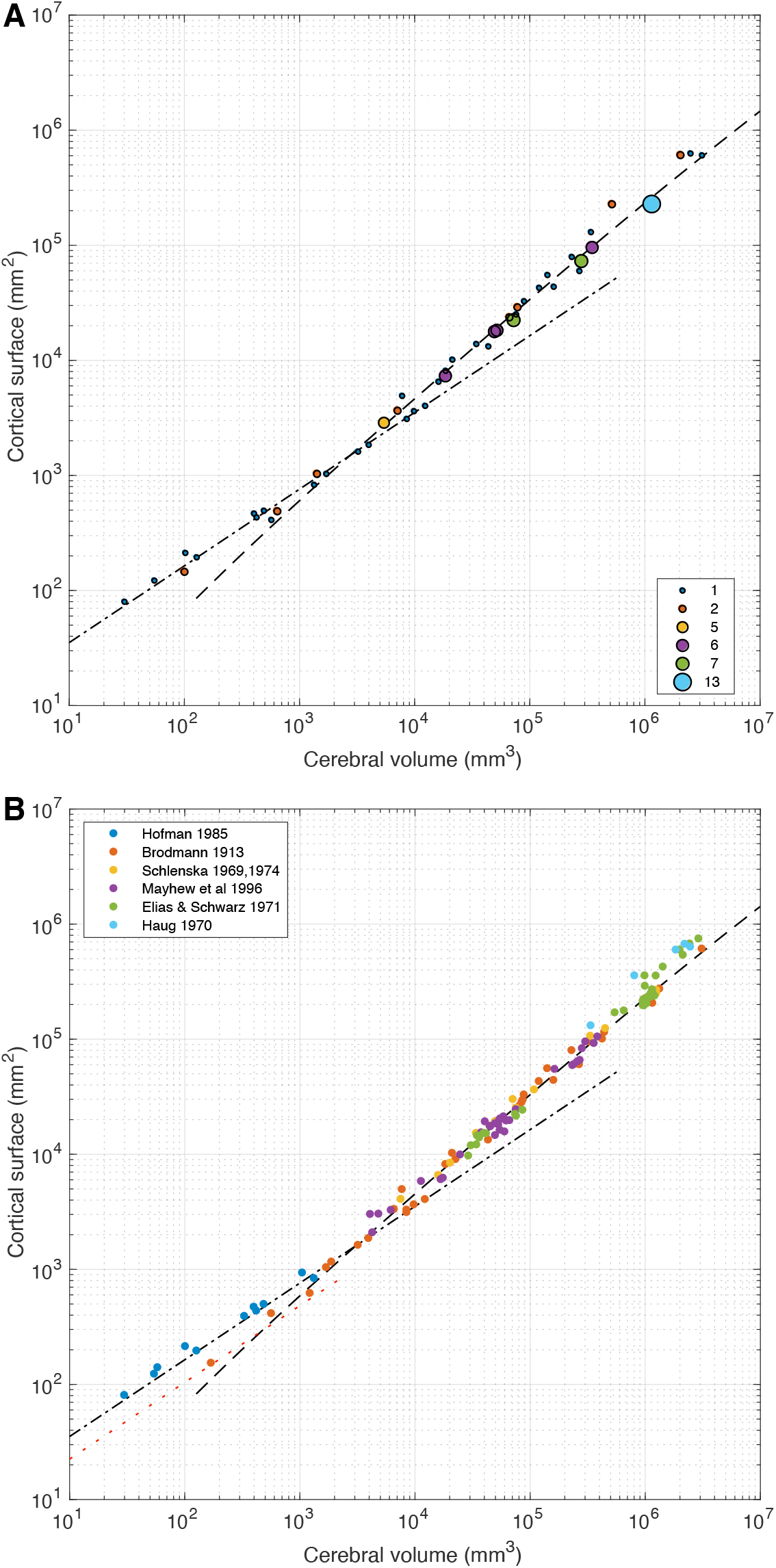
Supplemental Figure: Model for the cortical surface in relation to the cerebral volume (cf. Fig. 1). A. By no of specimen per species: the size of each icon represents the number of specimen included for each species. Humans are represented by the largest number, but note, that the MRI data of Toro et al (2008) was not included in the statistics. B. By reference. The individual specimen are each shown as a single dot. Note, that the surface area cannot be smaller than the surface area of a sphere of the same volume (i.e., the sphericity *s* > 1, see Fig. S1). This limit is shown by the red dotted line.

